# Neuroinflammation is an early pathological process in a mouse model of prodromal Parkinson’s disease

**DOI:** 10.64898/2026.10.29.762888

**Authors:** Gabriela Mercado, Lucas Stetzik, Emily Schulz, Lindsay Meyerdirk, Allison Lindquist, Lorna Cohen, Martha L. Escobar Galvis, Jennifer A. Steiner, Darren Moore, Daniel W. Wesson, Patrik Brundin

**Author notes:** Roche Pharma Research and Early Development (pRED) Neuroscience & Rare Diseases, Roche Innovation Center Basel, F. Hoffmann-La Roche Ltd., Basel, Switzerland.

## Abstract

Approximately 90% of Parkinson’s disease (PD) patients exhibit olfactory deficits, which often precedes motor symptoms by years, and are a robust predictor for PD and subsequent cognitive decline. Pathological changes in the PD olfactory system, particularly α-synuclein (αSyn) pathology in the olfactory bulb (OB) and severe involvement of the anterior olfactory nucleus (AON), are distinct from normal aging. However, the causal link between αSyn aggregation and functional impairment, potentially involving cytotoxicity and neuroinflammation, remains elusive. To investigate these early pathological events, we utilized a prodromal PD model that we have previously characterized, involving the unilateral OB injection of αSyn pre-formed fibrils (PFF) in wild-type mice. This triggers the transneuronal spread of pathogenic αSyn, and olfactory dysfunction starting at 1-month post-injections. Here we report on neuropathological and inflammatory events 1–3 month following injection of PFFs into the OB. Following PFF injection, αSyn pSer129 pathology progressively accumulated bilaterally in the AON and ipsilateral in the piriform cortex (PCX) over three months. Stereological quantification revealed no significant neuronal loss in the AON or PCX during this 1–3 month interval, indicating that olfactory impairment precedes overt neurodegeneration. Instead, we identified the upregulation of pro-inflammatory genes (*Il- 1β* and *Il-6*), significant microglial reactivity, and sustained astrogliosis within olfactory structures. These findings identify neuroinflammation as a key early mediator linking αSyn pathology to olfactory dysfunction in the absence of neuronal loss. This suggests a critical window for therapeutic intervention and highlights neuroinflammatory markers as potential biomarkers for the prodromal phase of PD.

**Highlights:**

1. **Neuroinflammation Precedes Neuron Loss.** Inflammation is an early pathological process that is detected before any significant loss of NeuN-positive neurons in olfactory structures.
2. **Progressive αSyn pSer129 spreading.** Pathological αSyn pSer129 progressively spreads transneuronally from the injection site in the OB to secondary olfactory structures (AON and PCX) within 1 to 3 months.
3. **Early Microglial Reactivity.** The spreading of αSyn pSer129 pathology rapidly induces microglial reactivity, which is prominent at the 1- and 2-month time points.
4. **Persistent Astrogliosis.** Astrogliosis is also an early and persistent component of the neuroinflammatory response, starting at 1 month and remaining detectable through 3 months, suggesting a sustained inflammatory environment.

## Introduction

Approximately 90% of people living with Parkinson’s disease (PD) experience olfactory deficits (Doty, 2012; Wu et al., 2011) which typically manifest 4-10 years before the onset of motor symptoms (Ross et al., 2008; Wu et al., 2011). Olfactory impairment is a strong predictor for both PD development (Lerche et al., 2014; Ross et al., 2008) and subsequent cognitive decline (Fullard et al., 2016; Morley et al., 2011; Schrag et al., 2017). Consequently, olfactory deficits are highly relevant for the early detection of PD and for stratifying patients in clinical trials targeting cognitive decline. Despite this importance, the neurobiological mechanisms of olfactory impairments in PD remain poorly understood.

Pathological changes in the olfactory system in PD differ from those observed during normal aging (Morley et al., 2011) and in other diseases like Alzheimer’s disease (Fusetti et al., 2010; Rey et al., 2018b). αSyn pathology in the olfactory bulb (OB) discriminates PD from non-PD elderly controls with high sensitivity (95%) and specificity (91%) (Beach et al., 2009b). Immunohistochemical, volumetric, and functional neuroimaging analyses suggest that loss of olfaction in PD is not due to damage of the olfactory epithelium but rather to alterations in central olfactory structures (Hummel et al., 2010). Lewy pathology in PD, characterized by αSyn aggregates, is present in mitral cells and in the inner plexiform layer of the OB, as well as in several cell subtypes in downstream olfactory structures (Beach et al., 2009a, 2009c; Ubeda-Bañon et al., 2010). Among these pathologically-burdened downstream structures, the anterior olfactory nucleus (AON) is the most affected (Ubeda-Bañon et al., 2020, 2017). However, the precise mechanism by which αSyn aggregates lead to impaired olfaction is largely unknown.

αSyn-related cytotoxicity (Wales et al., 2013) and neuroinflammation (Allen Reish and Standaert, 2015; Doorn et al., 2014b, 2014a; Moehle and West, 2015) are considered crucial and possibly synergistic molecular mechanisms implicated in PD pathogenesis. While αSyn pathology in postmortem olfactory structures is well documented, it is unclear how the presence of αSyn aggregates and neuroinflammation contribute to the functional deficits, and whether loss of neurons is required before olfactory dysfunction becomes apparent. Defining these underlying neurobiological mechanisms is essential for identifying novel therapeutic targets.

To investigate the contribution of αSyn neuropathology to prodromal olfactory deficits, we previously developed and characterized a mouse model of αSyn pathology in the olfactory system that exhibits olfactory deficits and mimics the pre-motor stages of PD. This model is based on the unilateral injection of recombinant αSyn preformed fibrils (PFF) directly into the OB of wild-type mice. We found that the injection of PFFs triggers the transneuronal transfer and seeding of pathogenic αSyn, spreading progressively to structures situated multiple synapses away, resulting in severe olfactory perceptual dysfunction. Mild olfactory deficits are detectable as early as 1 month post-injection, worsen over the subsequent two months, and then remain relatively stable for over 12 months (Rey et al., 2018a, 2016).

This early functional deterioration correlates with the progressive accumulation of αSyn pathology in the brain olfactory structures innervated by OB projection neurons, including the AON and the anterior and posterior piriform cortex (PCX). While no detectable mitral cell loss occurs in the OB even 18 months post-injections, cell loss has been reported in the AON at 6 and 18 months (Rey et al., 2018a). Crucially, no studies have investigated these OB targeted structures at early time points, leaving a gap in information regarding the process linking αSyn pathology and the progressive phase of olfactory dysfunction.

In this study, we concentrate on progressive phase of the functional impairment by investigating αSyn pathological spreading from the OB during early time points (1 to 3 months) post-PFF injection, with the aim of determining whether neurodegeneration or neuroinflammation are present during the early stages. We assessed neuronal loss via stereological quantifications in the AON and PCX. We also analyzed changes in gene expression for cellular pathways including synaptic function, inflammatory and stress pathways. Furthermore, we employed deep learning AI models for the easy visualization and extensive quantification of progressive αSyn pSer129 pathology, microglial reactivity and astrogliosis in olfactory structures. Collectively, these analyses demonstrate that inflammation is an early pathological process that precedes overt neuronal loss in this prodromal PD model.

## Methods

### Animals

We utilized 7- to 8-week-old C57BL6J wild-type mice purchased from Jackson’s Laboratory (Strain:000664). For all histological analysis a cohort of 16 animals per group (8 males and 8 females) were used per treatment (PBS and αSyn PFF) and per time point (1-, 2-, and 3-months). For quantitative real time PCR (qPCR) analysis, a different cohort of animals was used, with 8 animals per group (4 males and 4 females) per treatment (PBS and αSyn PFF) and per time point (1- and 3-months). Mice were housed in the vivarium at the Van Andel Institute, with a maximum of 4 animals per cage, under a 12- hour light-/dark cycle with *ad libitum* access to water and food. All animal housing and procedures were carried out in accordance with the Guide for the Care and Use of Laboratory Animals (United States National Institutes of Health) and were approved by the Van Andel Institute’s Institutional Animal Care and Use Committee (AUP 19-12-031).

### Intracerebral stereotaxic injections

Mouse αSyn PFF were produced as described in (Volpicelli-Daley, Luk et al. 2014) and kindly provided by Dr. Kelvin Luk at University of Pennsylvania Perelman School of Medicine, USA. Mice were anesthetized with isoflurane and unilaterally injected into the right OB (coordinates from bregma: AP: + 5.4 mm; ML: +/− 0.75 mm and DV: −1.0 mm from dura) with 0.8 µL of PFF [5 µg/µL]; or 0.8 µL of PBS (phosphate buffered saline) as control. Injections were performed at a rate of 0.2 µL/min using a glass capillary attached to a 10 µL Hamilton syringe. After injection, the capillary was left in place for five minutes before slow removal. Prior to incision, the animals received a local anesthetic injection at the wound margin (0.03 mL of 0.5% Ropivacaine; Henry Schein, USA). Following surgery, mice received 0.5 mL saline subcutaneous (s.c.) for hydration, and 0.04 mL Buprenex (Henry Schein,) s.c. for analgesia.

### Tissue collection and quantitative real-time PCR

Animals were euthanized at 1 or 3 months after stereotaxic injections with sodium pentobarbital (130 mg/kg; Sigma) and transcardially perfused with PBS to remove blood. Brains were isolated, and the OB, AON and PCX were rapidly dissected on ice using a mouse brain matrix. Isolated samples were weighed, snap frozen, and stored at –80°C for later analysis. Tissue was homogenized in PBS followed by sonication. Total RNA was isolated from the injected OB, AON and PCX using the RNeasy mini kit (QIAGEN). Quantitative real time PCR (qPCR) was performed in technical duplicates on a StepOne™ Real-Time PCR System using TaqMan™ RNA-to-CT™ 1-Step Kit (Applied Biosystems) and TaqMan™ Gene Expression Assay (FAM) probes for 10 targets listed in **Supplementary Table 1**. Gene expression levels were determined using the Relative Quantification (RQ) app from Thermo Fisher Connect^TM^. *Gapdh* was used as specific endogenous control on each sample, and data was graphed using the biological group results relative to the time matched PBS-injected control group (n=8 biological replicates for each of the 4 experimental groups).

### Immunohistological staining

Animals were anesthetized with sodium pentobarbital (130 mg/kg; Sigma Aldrich) for tissue collection at the indicated time points. Brains were isolated after transcardial perfusion with saline and fixation with 4% PFA. They were then post-fixed overnight with 4% PFA and then stored at 4°C in 30% sucrose in PBS until sectioning. Brains were frozen, and coronal sections (40 µm) were cut on a sliding microtome (Leica) and collected as serial tissue sections. For immunohistochemical (IHC) analysis, a series of coronal free-floating tissue sections (every 6^th^ section or 240 µm intervals) were used. Samples were quenched with 3% H2O2 at room temperature (RT) for 30 min and blocked with 10% normal goat serum, 0.4% bovine serum albumin, and 0.1% triton-X in PBS for 2 hours. They were then incubated with primary antibodies listed in **Supplementary Table 2** at 4°C overnight. The following day, samples were washed three times with 0.1% Triton-X in PBS and incubated with the biotinylated secondary antibody (**Supplementary Table 2**) for 2 hours at RT, triple washed again and treated with Vectastain Elite ABC HRP kit (Vector Laboratories, PK-4000). Antigen detection was performed with the Vector DAB substrate kit (Vector Laboratories, SK-4100). After mounting and dehydration, slides were coverslipped with Cytoseal 60 mounting medium (Thermo Fisher Scientific). Fluoro-Jade C staining was performed following manufacturer’s instructions (Millipore, AG325-30MG). For immunofluorescence (IF) analysis, when indicated antigen retrieval was performed with the universal HIER antigen retrieval reagent (Abcam, ab208572) for 30 minutes in a steam cooker at 95-100°C. Tissue sections were blocked with 10% donkey normal serum (Millipore) and 4% bovine serum albumin in 0.1% Triton-X in PBS for 2 hours at RT. Samples were then incubated with primary antibodies listed in **Supplementary Table 3**. The next day, slides were triple washed with 0.1% Triton-X in PBS and incubated with secondary antibody (**Supplementary Table 3**) for two hours at RT after which the sections were washed again with 0.1% Triton-X in PBS. Slides were sealed with Vectashield Antifade Mounting Medium with or without DAPI (Vector Laboratories).

### Image acquisition for immunofluorescence staining

The sections were viewed under a Nikon Eclipse Ni-U microscope (Nikon). Brain tissue sections were imaged on a Zeiss LSM 880 equipped with an AiryScan detector in Fast mode and a Plan-Apochromat 63x oil immersion lens (N.A. 1.4), in Zen Black (ver. 2.3) software. Images were Z-stacked at a total depth of 15 µm in 1 µm intervals, and in a 3x3 tile. Post-acquisition, all images underwent AiryScan processing and tile-stitching.

### Stereology

NeuN-positive cells were quantified by unbiased sampling and blinded stereology, employing a section evaluation interval of 6 and the optical fractionator probe in Stereo Investigator software (MBF Bioscience). Contours of the AON, aPCX and pPCX were drawn at 2.5x magnification, according to the Allen Mouse Brain Atlas (Allen Institute). Quantifications were made at 100x magnification using a 200 µm counting frame. The dissector height was manually measured on each frame with a guard zone of 3 µm, the grid was set to 200 × 200 and the frame to 40 × 40 respectively. The Gunderson estimated m = 1 error was less than 0.1.

### Automated histopathological assessment

Z-stacks of mounted IHC-stained tissue sections were captured at 20x magnification using a whole slide scanner (Zeiss, Axioscan Z1) at a 0.22 μm/pixel resolution. Extended Depth Focus (EDF) was used to collapse the z-stacks into 2D images as they were collected. Tissue thickness was set to 20 μm, z-stacks were collected at 3 μm intervals, 3 μm optical slices were chosen based upon optimal contrast. The digitized whole slide images were uploaded to Aiforia® image processing and management platform (Aiforia® Cloud, Aiforia Inc.) for analysis with deep learning convolutional neural networks (CNNs) and supervised learning. Regions of interest (ROI) in the injected and non-injected OB, AON, anterior PCX (aPCX) and posterior PCX (pPCX) were outlined in the coronal serial tissue sections using as reference the Allen Mouse Brain Atlas (Allen Institute) and tools in the Aiforia® Cloud platform. Quantifications were performed in all the outlined ROI using specific models for different IHC staining. Quantification of phosphorylated αSyn (pSer129) area % and counts per mm^2^ was performed in the Aiforia® platform using our previously reported deep learning assisted model (Bergkvist et al., 2021; Stetzik et al., 2024). The automated assessment of microglia morphology was performed on anti-Iba1 IHC-stained brain tissue section images as previously described (Mercado et al., 2024; Stetzik et al., 2022). To quantify astrogliosis, we developed a new model using images from anti-GFAP IHC-stained brain tissue and similar methodology to those previously described (Stetzik et al., 2022) (**Supplementary Figure 1**). In brief, a multi-layered CNN was trained on the most diverse and representative images across the datasets using tools in the Aiforia® Cloud platform. The first layer was annotated using semantic segmentation to detect total tissue area. The second segmentation layer to detect total GFAP positive area. The third was a GFAP-positive soma counter layer. For each layer, features that were considered artifacts were annotated as background and used as additional input during model training iterations.

### Statistical analysis

For the real time PCR data, p-values were calculated in the RQ app from Thermo Fisher Connect^TM^ using two-tailed unequal variance t-test and the Benjamini-Hochberg correction to control false discovery rate for multiple comparisons. For the histological data sets, statistical analyses were conducted using GraphPad Prism 10 software. Each data point used represents one animal average of the selected ROIs in serial tissue section every 6^th^ section or 240 µm intervals encompassing the entire brain arears (OB, AON, aPCX and pPCX). The type of analysis with post hoc correction for multiple comparison is indicated in the legend of each figure. Statistical levels were set at *p <0.05, **p<0.01, ***p<0.001.

## Results

### αSyn pSer129 pathology progressively spreads through olfactory structures after αSyn PFF injection in the OB

We used a deep learning AI algorithm for quantification and easy visualization of the progressive spreading of αSyn pSer129 pathology in olfactory structures 1, 2 and 3 months after PBS or αSyn PFF injections into the right OB of wild-type mice (**Figure 1A**). Our sensitive unbiased method allows the quantification of αSyn pSer129 pathology, measured as the percentage of the area positive for the pathological marker and the total positive counts per brain area. Both metrics showed highly significant main effects for treatment and hemisphere, with a strong treatment x hemisphere interaction, confirming that the effect of αSyn PFF injections are confined to the injected hemisphere (shown in red, **Figure B**).

**Figure 1.**
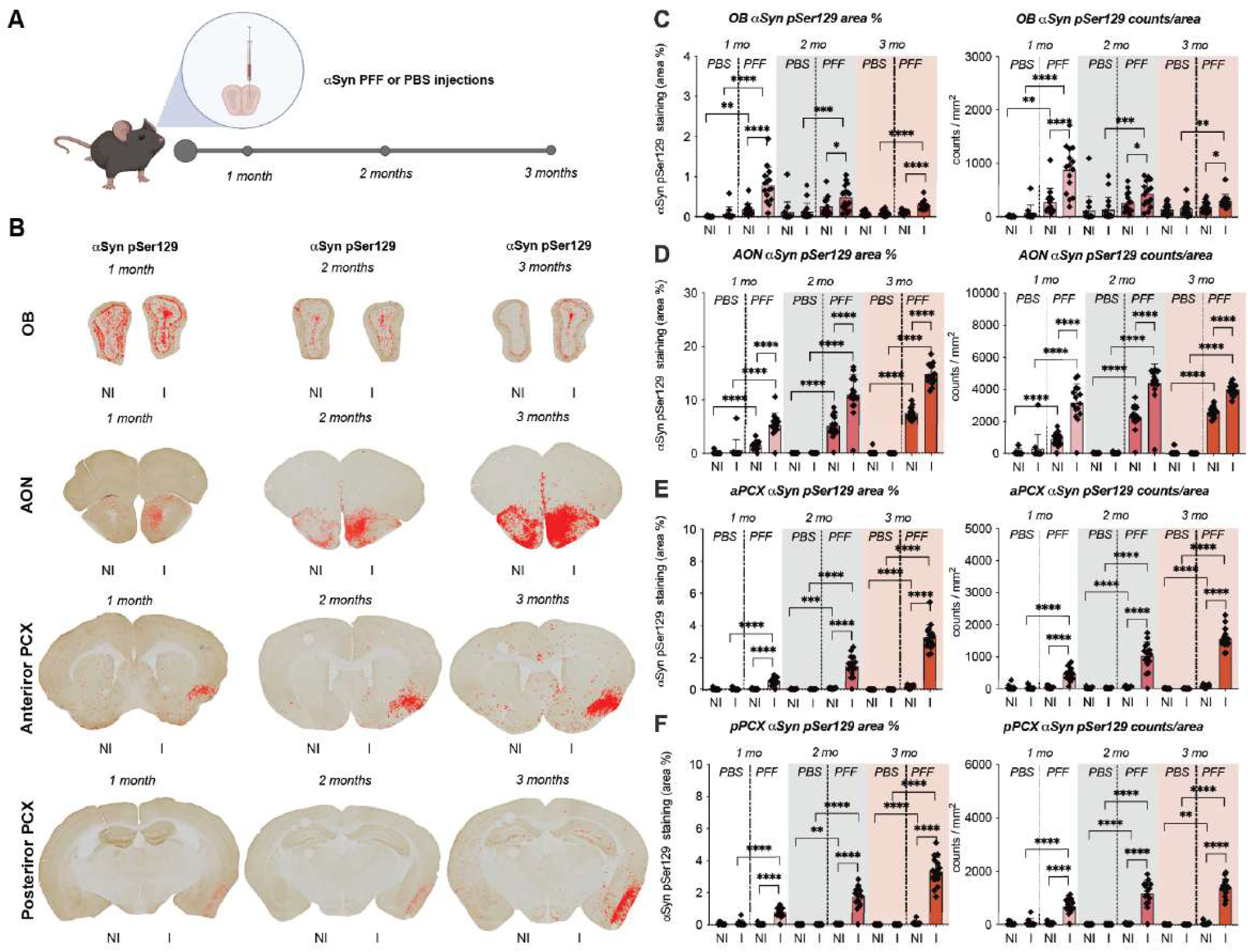
Spatiotemporal spreading of αSyn pSer129 after PFF injection in the OB. **(A)** Wild- type mice were unilateral injected in the OB with αSyn PFF or PBS, and brains were collected for analysis at 1, 2 and 3 months post-injection. Anti-αSyn pSer129 DAB-stained serial tissue sections images were analyzed using a deep learning algorithm. **(B)** Representative images of αSyn pSer129 immunoreactivity visualization (in red) in brain olfactory structures from αSyn PFF injected animals. Non-injected (NI) and injected hemispheres (I), respectively. Quantification of αSyn pathology, calculated as the percentage of αSyn pSer129 positive immunoreactivity area of the total area (left panels) and as the number of αSyn pSer129 positive counts per area (right panels), is shown for the OB **(C)**, AON **(D)**, aPCX **(E)** and pPCX **(F)** at 1, 2 and 3 months (mo) after injections. Each data point represents the average per animal (n= 16 per group). Data was analyzed using three-way ANOVA (Factors: Time, Treatment, and Hemisphere). Post-hoc comparisons between injected and non-injected hemispheres, and treatment groups were performed using t-tests with a Bonferroni correction for multiple comparisons across time points.

As expected, specific post-hoc comparison for the injected *versus* non-injected side confirmed significantly higher pathology on the injected side across all regions, starting at 1 month. While the PBS-injected *versus* PFF-injected comparison highlights an effective PFF seeding effect throughout the olfactory system (**Figure 1C-F**). In line with our previous work (Rey et al., 2018a and 2016), αSyn pSer129 pathology exhibited a robust, time-dependent progression. While early pathology is concentrated at the injection site (OB) and AON, it significantly spreads to the cortical regions (aPCX and pPCX) by the 2–3 month marks (**Figure 1C-F**). Our results also highlight the AON as the most affected area, with progressive spreading to the contralateral side (**Figure 1D**). Overall, these analyses show that the αSyn PFF treatment was effective in seeding pathology, and the injected side consistently shows significantly higher pathology than the non-injected side.

A separate analysis including sex as a variable, showed that biological sex did not play a primary main effect, suggesting that in our model the pathology developed similarly in males and females. However, the post-hoc comparison revealed discrete sex-driven differences in the contralateral spreading of αSyn pSer129 pathology at the earliest time point of 1-month post-injections, females display an increase contralateral pathology in the OB, while males presented increased contralateral pathology in the aPCX (**Supplementary Figure 2A**). Although a similar model involving αSyn PFF injections in the AON previously reported higher ipsilateral αSyn pSer129 pathology in male’s mice olfactory structures 3 months post-injections (Mason et al., 2016), our quantifications do not support a significant sex-driven effect (**Supplementary Figure 2B**).

### αSyn pSer129 pathology spreading does not induce neuronal loss in olfactory structures early after PFF injection in the OB

We have previously reported that in this model of αSyn PFF injection into the OB, animals do not display neuronal loss in the OB measured 1, 3, 6 and 18 months post-injection, but that a significant cell loss is observed in the AON 6 and 18 months post-injection (Rey et al., 2018a, 2016). However, no data has been reported for the AON at earlier time points or for any other cortical area at any time point post-injection. To determine if the αSyn pathological spreading correlates with neuronal cell loss in olfactory structures at early time points, we performed stereological quantifications of cells positive for the neuronal nuclear marker (NeuN) in the AON, aPCX and pPCX 1, 2 and 3 months after PBS or αSyn PFF injections (**Figure 2A**). While there were fluctuations over time, the data analysis did not show a significant main effect of treatment or side on the total NeuN counts, suggesting that at these points (up to 3 months) the αSyn pathology present without causing neuronal loss in these regions detectable by NeuN stereological counts (**Figure 2B, C, D**). Across all regions, we did not observe that sex significantly alters the trajectory of the αSyn PFF-induced neurotoxic cascade (**Supplementary Figure 3**). We also tested for cell death markers in olfactory structures and found only discrete positive signals in the OB for cleaved Caspase 3 in the olfactory nerve layer and Fluoro-Jade C in the internal plexiform layer, both consistent with the stereotaxic needle track, and both markers being observed independent of the treatment (**Supplementary Figure 4**). Our data indicate that NeuN counts remain stable across all time points and treatment groups, suggesting that while αSyn pathology is widespread and progressive at 3 months, it has not yet reached the threshold for significant neuronal death in the AON and PCX.

**Figure 2.**
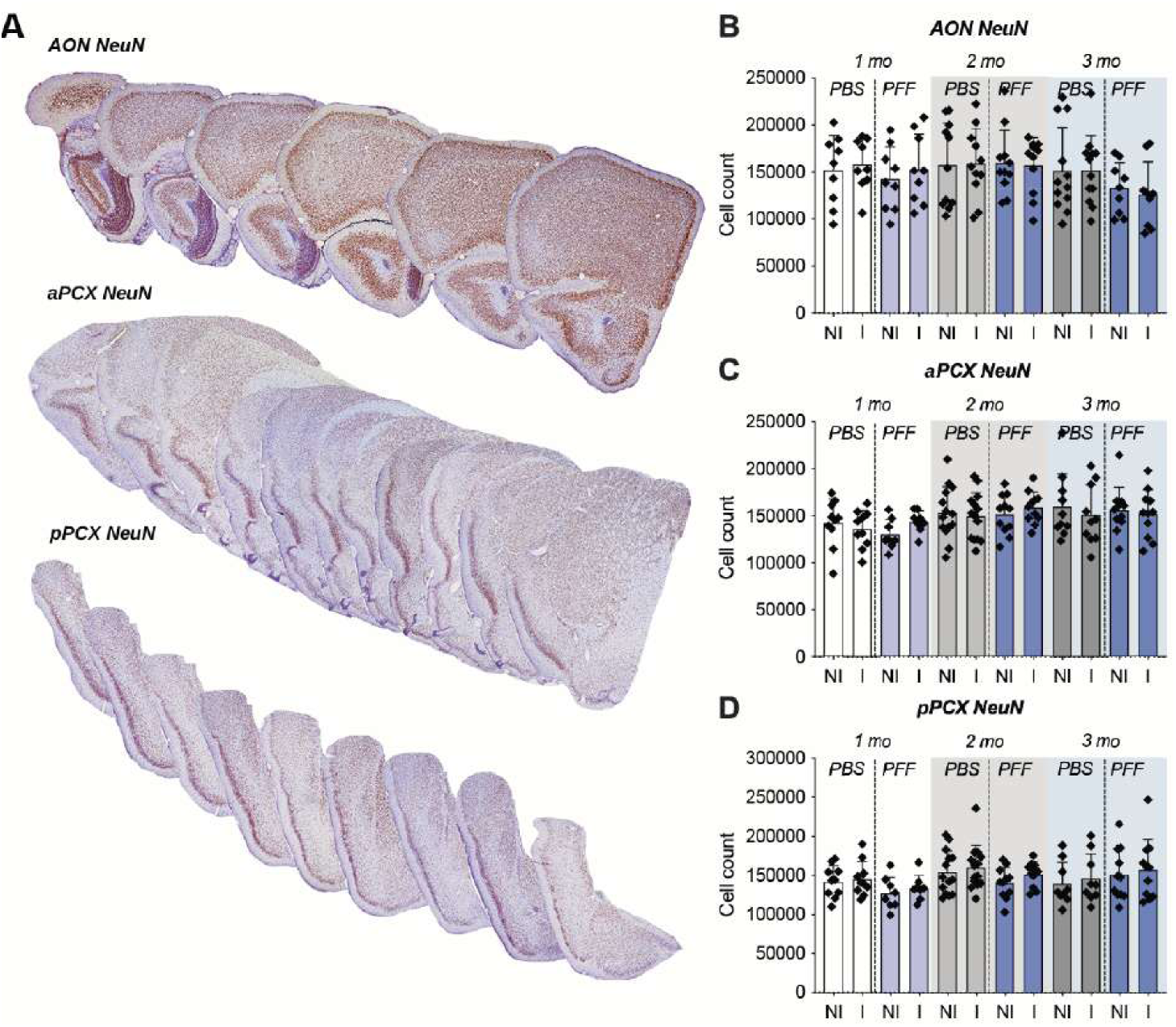
αSyn pSer129 pathology spreading does not induce NeuN positive cell loss in olfactory structures. Stereological quantifications of NeuN positive cells in brain olfactory structures 1, 2 and 3 months after αSyn PFF or PBS injections into the OB of wild-type mice. **(A)** Representative images of anti-NeuN DAB-stained serial tissue sections from the AON, anterior PCX (aPCX) and posterior PCX (pPCX). Total cell counts for the AON **(B)**, aPCX **(C)** and pPCX **(D)** at 1, 2 and 3 months (mo) after injections. Non-injected (NI) and injected hemispheres (I), respectively. Each data point represents the average per animal (n= 16 per group). Data analyzed using three-way ANOVA (Factors: Time, Treatment, and Hemisphere).

It has been previously described, in both PD patients and mouse models, that αSyn pathology is frequently distributed in glutamatergic mitral cells, inhibitory interneurons, and interneurons expressing calcium binding proteins, while dopamine and somatostatin positive neurons are partially or rarely affected (Beach et al., 2009a, 2009c; Ubeda-Bañon et al., 2010). To determine if the αSyn pSer129 pathological marker is distributed in specific neuronal subpopulation(s) in our model, we used co-immunofluorescent staining with antibodies specific for inhibitory neurons, excitatory neurons, different interneurons, and interneurons expressing calcium binding proteins (**Supplementary Table 3**) in PFF-injected animals at 3 months. These co-immunofluorescent staining experiments demonstrated that, as observed in postmortem tissue from PD patients, the αSyn pathology preferentially accumulates in mitral cells (Tbr2-positive), inhibitory cells (ChAT-positive) and calcium binding protein expressing interneurons (Calreticulin- and Parvalbumin-positive; **Supplementary Figure 5**). Although less abundant, we also observed some αSyn pSer129 pathology co-distributed with the markers vGlut1, somatostatin and cholecystokinin B receptor (Cckbr) while no observable αSyn pSer129 pathology was found in TH- positive (dopaminergic) interneurons (**Supplementary Figure 5**). These observations show that, as described in other animal models and human studies (Beach et al., 2009a, 2009c; Ubeda-Bañon et al., 2010), there appears to be a differential susceptibility of inhibitory interneurons but not a unique neuronal subpopulation affected by αSyn pathology.

### Increased levels of pro-inflammatory cytokines in olfactory structures after αSyn PFF injection in the OB

Since we did not observe substantial neuronal loss in olfactory structures that could account for the previously reported functional impairment induced by αSyn PFF injection into the OB, we performed a gene expression analysis of bulk tissue from olfactory structures to obtain clues about the underlying biological process. We used validated probes to measure the expression levels of markers of a wide range of cellular pathways, including synaptic function (*Snca* and *Dlg4*), apoptosis (*Chop*), inflammation (*Nlrp3, Il-6* and *Il-1β*), astrogliosis *(Gfap*), endoplasmic reticulum (ER) stress (*Hspa5*), lysosomal vesicles (*Lamp1*) and mitochondrial function (*Nox2*) (**Supplementary Table 1**).

Quantification of relative mRNA levels (RQ) were performed for the PBS- or PFF- injected OB, AON and PCX at 1 and 3 months after injections, using *Gapdh* as endogenous control (**Figure 3**). Most of the significant changes in gene expression were observed in the OB, including increased ER stress marker Bip (*Hspa5*) at 1 month, and increased levels of the pro-inflammatory marker *Il-1β* at 1 and 3 months after PFF injections (**Figure 3A**). In the other analyzed brain regions (AON and PCX), we observed higher inter-animal variability than in the OB and we only observed significative increase of *Il- 6* in the AON 1-month post PFF-injection (**Figure 3B**); however, increased levels of *Il-1β* and *Il-6* were observed across brain regions and time points (**Figure 3**). These results suggest that inflammation could be an early pathological process that correlates with αSyn pathology in this mouse model of prodromal PD.

**Figure 3.**
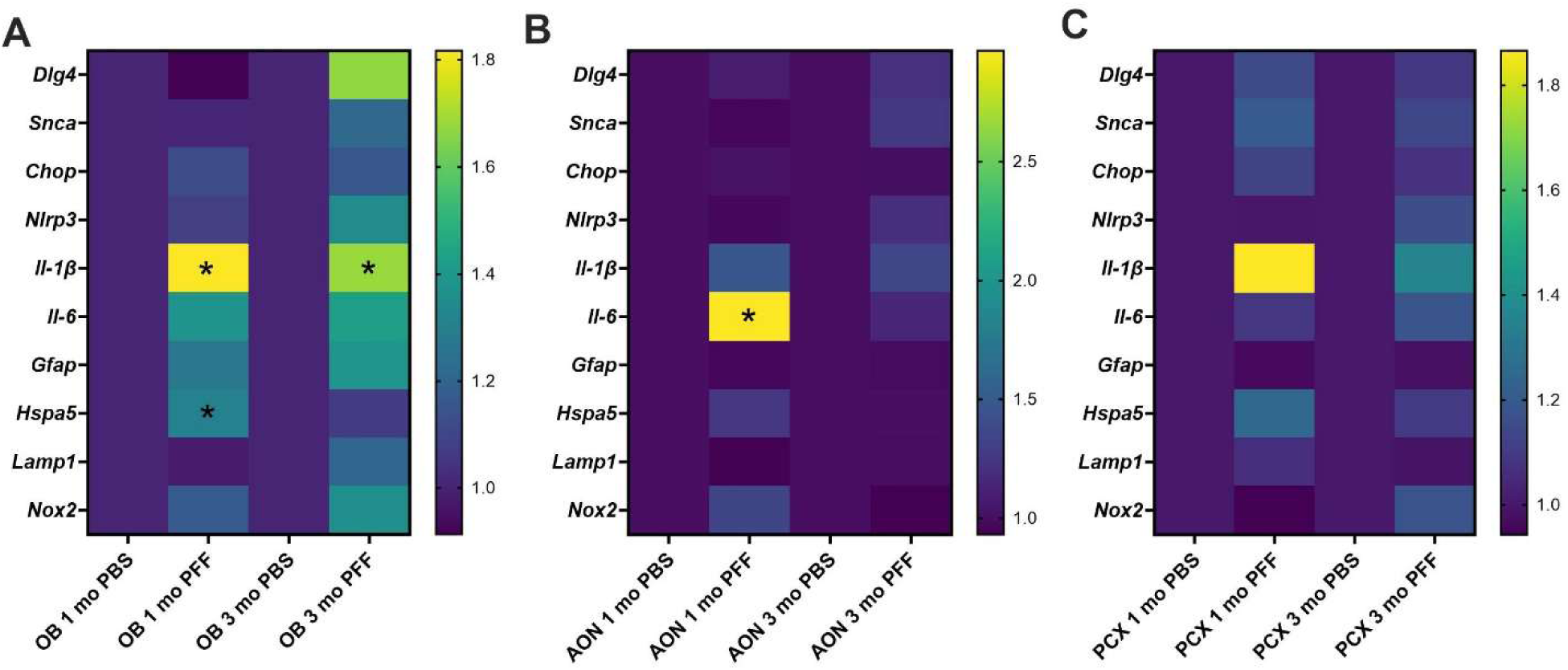
Gene expression levels in olfactory structures by qPCR after ⍺Syn PFF injections in the OB. Relative mRNA levels of *Snca, Dlg4, Chop*, *Nlrp3*, *Il-1β*, *Il-6*, *Gfap*, *Hspa5*, *Lamp1* and *Nox2* measured by real-time quantitative PCR (qPRC) in the **(A)** OB, **(B)** AON and **(C)** PCX from αSyn PFF- and PBS-injected animals 1- and 3-months after injections. *Gapdh* was used as endogenous control and relative quantification (RQ) are expressed as fold of change compared to time matched PBS control group (n=8 per group, half male and half female; p > 0.05 with unequal variance two-tailed t-test and the Benjamini-Hochberg correction for multiple comparisons).

### Increased microglial reactivity after αSyn PFF injection into the OB

To characterize the neuroinflammatory response induced by αSyn PFF injections into the OB, we used deep learning assisted models to quantify reactive microglia signatures across the olfactory structures. Various pathological inputs can induce microglial reactivity, which is characterized by changes in their functional state, typically reflected by morphological changes (Area:Perimeter ratio changes) and microgliosis (an increase in the number of cells per area). We used anti-Iba1 DAB-stained histological images and a deep learning AI algorithm to quantify microglial morphology and microgliosis in olfactory structures (OB, AON, aPCX and pPCX) 1, 2 and 3 months after PBS (**Figure 4A**) or PFF injections (**Figure 4B**). Our unbiased method for cross-regional quantification of microglial activation showed significant effects of time and treatment, indicating a progressive response (**Figure 4C**). Iba1 counts showed significant main effects of time and hemisphere, suggesting an inflammatory response wherein microglia proliferate and migrate to the areas of active pathology (**Figure 4C**).

**Figure 4.**
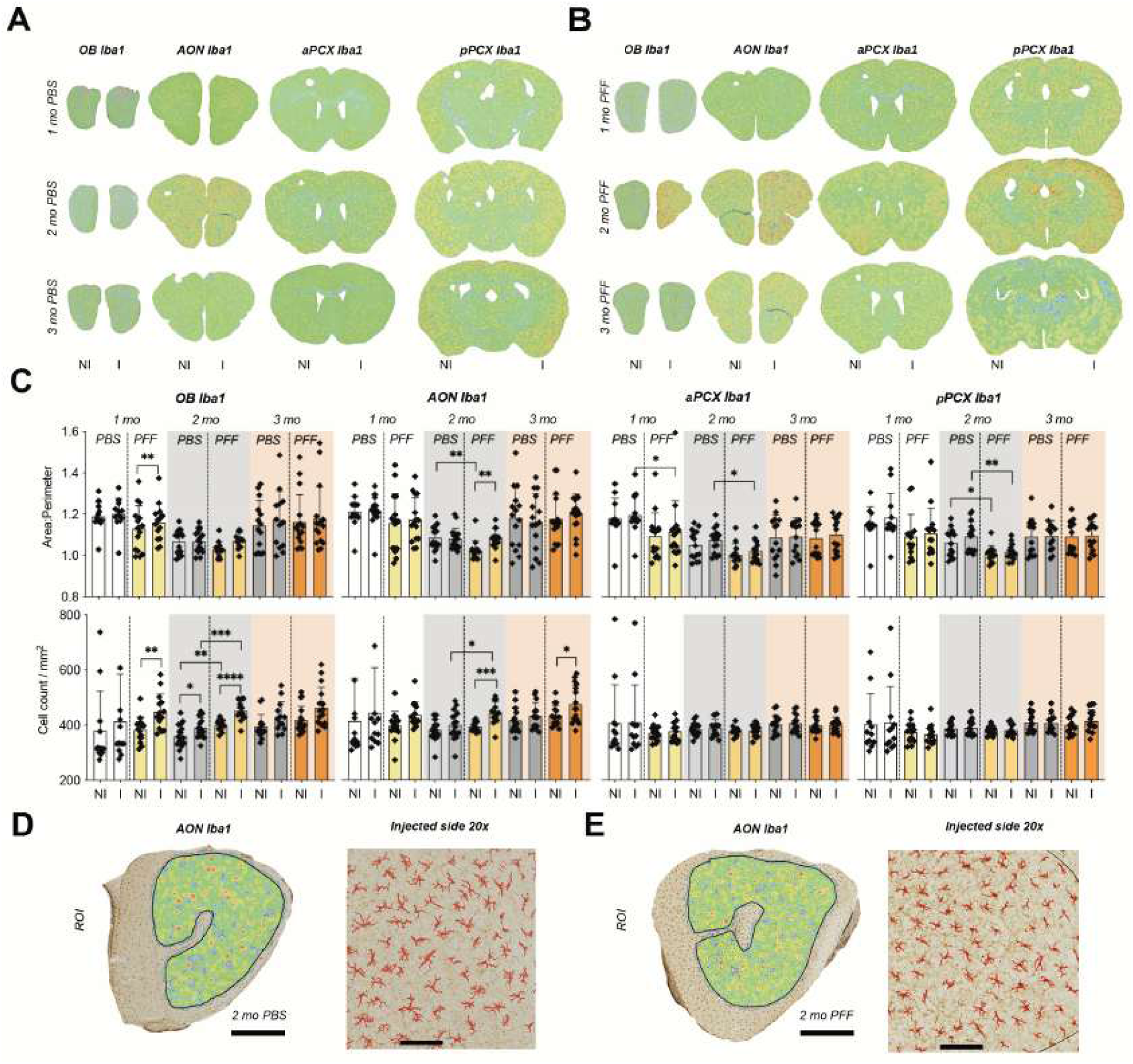
Increased microglial reactivity after ⍺Syn PFF injection in the OB. Anti-Iba1 DAB- stained serial section images were analyzed using a deep learning algorithm. Representative images of Iba1-positive cells visualization (heat map) in the brain of **(A)** PBS- and **(B)** αSyn PFF- injected animals 1, 2 and 3 months after injections. Non-injected (NI) and injected hemispheres (I), respectively. **(C)** Quantification microglial morphology expressed as the area-to-perimeter ratio of Iba1-positive cells (upper panels) and microgliosis as the number of Iba1-positive cell counts per area (lower panels) in the OB, AON, aPCX and pPCX. Each data point represents the average per animal (n= 16 per group). Data was analyzed using three-way ANOVA (Factors: Time, Treatment, and Hemisphere). Post-hoc comparisons between injected and non-injected sides, and treatment groups were performed using t-tests with a Bonferroni correction for multiple comparisons across time points. Representative images of the AON with visualizations of Iba1- positive cells (left panels, scale bar 500 µm) and the 20x magnification of the injected hemisphere (right panels, scale bar 100 µm) of PBS **(D)** or αSyn PFF **(E)** 2 months after injections.

The specific comparison at each time point showed that αSyn PFF-induced changes in microglial morphology were significant at 1 month post-injection in the aPCX, and at 2 months post-injection in the AON, aPCX and pPCX. No significant morphological changes were found in the analyzed olfactory structures after 3 months (**Figure 4C, upper panels**). Changes in the number of Iba1-positive cells per area (microgliosis) were significant in the OB at 1 and 2 months post-injection and in the AON at 2 and 3 months (**Figure 4C, lower panels**). Interestingly, some reactive microglia changes were present bilaterally, in the injected and non-injected hemispheres, such as the Area:Perimeter ratio in the pPCX and microgliosis in the OB 2 months post-injections (**Figure 4C**). We selected the AON 2 months after injections as a representative area to show high resolution images and the performance of the Iba1 deep learning assisted model for PBS- and PFF-injected animals (**Figure 4D and E**, respectively).

While sex did not show a main effect, we did identify sex-specific differences in the microglial morphology. Females showed increased microglial reactivity at the early 1-month post-injection time point, both ipsilateral and contralateral to the injection site, regardless of whether they were injected with PBS or PFF. In contrast, males only show increased microglial reactivity at 2 and 3 months post-injection in both hemispheres but only in PFF-injected animals (**Supplementary Figure 6**).

### Increased astrogliosis after αSyn PFF injection into the OB

To quantify astrogliosis in the same αSyn PFF spreading mouse model, we developed a deep learning assisted model that allow us to quantify the total area coverage of GFAP-positive immunoreactivity and the total number of GFAP-positive cells from anti-GFAP DAB-stained histological images (**Supplementary Figure1**) in olfactory structures (OB, AON, aPCX and pPCX) 1-, 2- and 3- months after PBS (**Figure 5A**) or PFF injections (**Figure 5B**). This new method for extensive cross-regional quantifications showed that reactive astrogliosis increased in response to PFF treatment over time, mirroring the spatial and temporal spread of αSyn pathology (**Figure 5C**).

**Figure 5.**
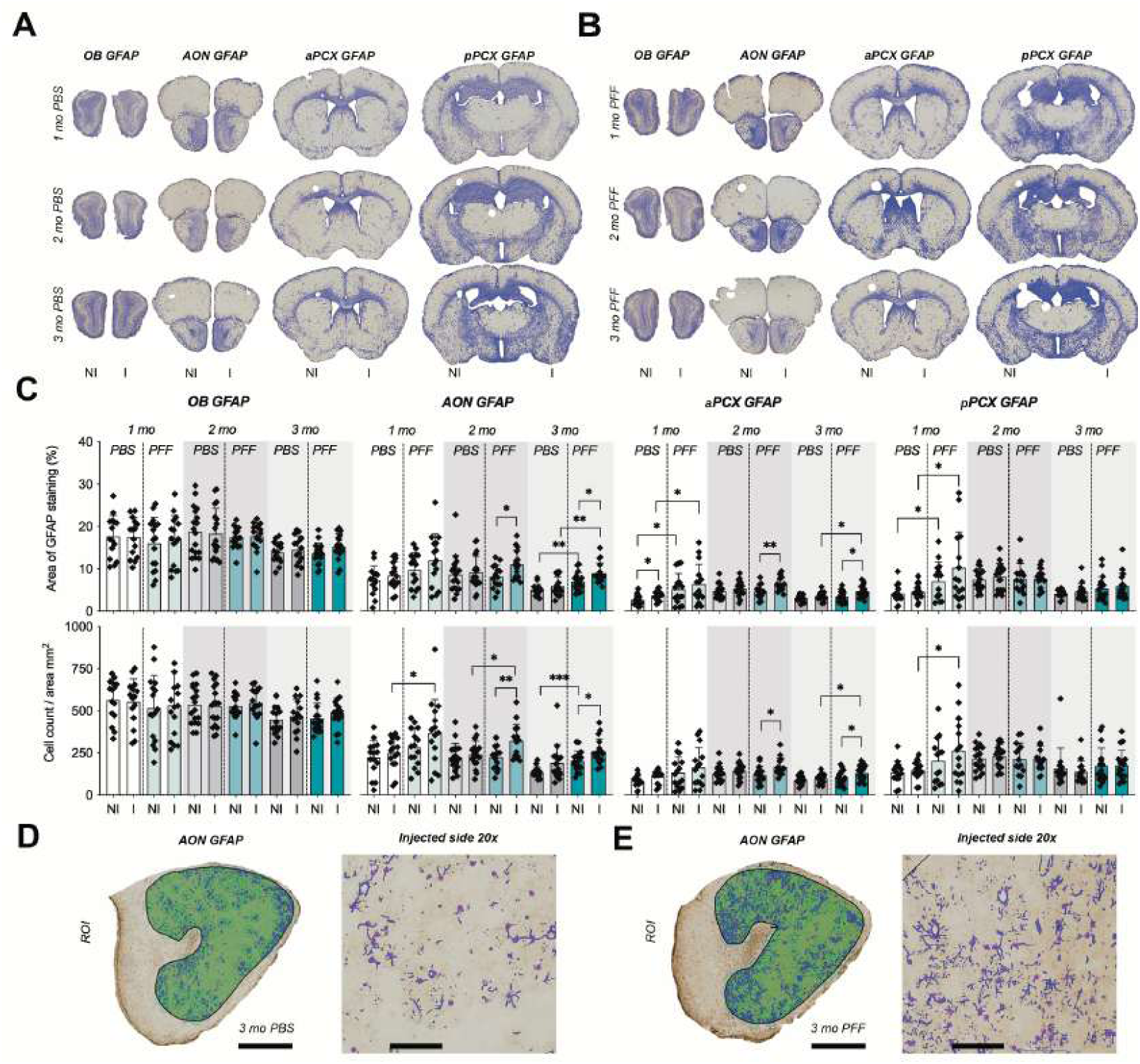
Increased astrogliosis in olfactory structures after ⍺Syn PFF injection into the OB. Anti-GFAP DAB-stained serial tissue section images were analyzed using a deep learning algorithm. Representative images of GFAP-positive area visualization (in blue) in the brain of **(A)** PBS- and **(B)** αSyn PFF-injected animals 1, 2 and 3 months after injections. Non-injected (NI) and injected hemispheres (I), respectively. **(C)** Quantification of astrogliosis, calculated as the percentage of GFAP-positive area of the total area (upper panels) and as the number of GFAP- positive soma per area (lower panels) of the OB, AON, aPCX and pPCX. Each data point represents the average per animal (n= 16 per group). Data was analyzed using three-way ANOVA (Factors: Time, Treatment, and Hemisphere). Post-hoc comparisons between injected and non- injected hemispheres, and treatment groups were performed using t-tests with a Bonferroni correction for multiple comparisons across time points. Representative images of the AON with GFAP-positive area and soma detection visualizations (left panels, scale bar 500 µm) of the injected hemisphere (right panels, scale bar 100 µm) of PBS **(D)** or αSyn PFF **(E)** 3 months after injections.

Specific comparison showed that αSyn PFF-induced changes in the area percentage covered by GFAP-positive immunoreactivity was significantly increased at 1-month post-injection in the aPCX and pPCX, and in the AON and aPCX 2 and 3 months after PFF treatment (**Figure 5C, upper panels**). For the GFAP-positive cells per area, we found increased astrogliosis in the AON and pPCX 1 month after treatment and in the AON and aPCX at 2 and 3 months (**Figure 5C, lower panels**). In contrast to microglial reactivity, where most significant differences were concentrated at the early time points (1 and 2 months post-injection), the astrogliosis was still persistent 3 months after PFF treatment. For the astrogliosis quantifications, we selected the AON at the 3-month time point as a representative area to show higher magnification visualizations and the performance of the GFAP deep learning assisted model for PBS- and PFF-injected animals (**Figure 5D and E**, respectively).

Sex did not show any significant main effect on astrogliosis after αSyn PFF spreading, however some specific comparisons showed sex-specific differences, with significant sex differences being more prominent ipsilateral to the injection site, and with male animals showing increased astrogliosis after both PBS and PFF injections (**Supplementary Figure 7**).

Given the persistence of astrogliosis after PFF injections and previous studies in PD patients’ samples describing astrogliosis (Mazumder et al., 2022; Pons-Espinal et al., 2024), we decided to define the presence of the αSyn pSer129 pathological marker in GFAP-positive astrocytes in the OB, AON and PCX. Using co-immunofluorescent staining, we observed that while there is an increase in GFAP immunoreactivity in the areas positive for αSyn pSer129, we did not observe this pathological marker in the GFAP-positive cells, suggesting that the functional effect of astrogliosis may be exerted through pro-inflammatory signaling pathways (**Supplementary Figure 8**).

We then performed a correlation analysis with the percentage area covered by αSyn pSer129- positive staining and the other histological markers across all regions and time points. We found a significant positive correlation with percentage area covered by GFAP staining (r = 0.295, p < 0.0001), indicating that a higher αSyn pathology is associated with increased astrogliosis. A significant positive correlation was also found with Iba1-positive counts (r = 0.356, p < 0.0001), suggesting that αSyn pathology triggers microglial proliferation/migration. Finally, no significant correlation was found with NeuN counts (r = –0.052, p = 0.32), reinforcing that pathology levels have not yet reached a threshold for significant neuronal death in this time frame.

## Discussion

The primary finding of this study is that neuroinflammation is an early pathological process in a mouse model of PD, which is parallel to the prodromal stage in humans. This robust response emerges and progresses before any significant neuronal loss is detectable in olfactory regions. This strong temporal correlation suggests that neuroinflammatory and glial responses, rather than the death of neurons, as potential drivers of the previously reported olfactory dysfunction in this model. Even though a causal link remains to be established, our observations position this model of prodromal PD as a useful tool to study the effects of anti-inflammatory interventions in the sensorial deficits and the progression towards neurodegeneration.

Our stereological quantifications of NeuN-positive counts revealed no significant differences across any of the olfactory brain regions examined, confirming the absence of overt neuronal loss despite high αSyn pathology burden in these regions (**Figure 2**). This observation contrasts with previous findings for later time points (6 and 18 months), where AON neuronal loss has been reported using the same methodology (Rey et al., 2018a, 2016). However, in neurodegenerative diseases including PD, synaptic loss frequently precedes somatic cell death and often correlates with functional decline. Investigating synaptic densities, levels of synaptic markers, or functional electrophysiological studies would be required to further confirm previous observations that support a model where synaptic dysfunction and neural network dysfunction, rather than neuronal death, underlies the initial functional decline (Kulkarni et al., 2022, 2020).

In our spatiotemporal analysis of αSyn pSer129 accumulation, the pathology originates in the injected OB and disseminates transneuronally to the AON and PCX, confirming the model’s fidelity to the anatomical spread of Lewy pathology described in human PD (**Figure 1**). The pathology, over time, was markedly bilateral in the AON but remained mostly ipsilateral in the PCX, suggesting differential susceptibility or connectivity patterns that govern the rate and extent of transneuronal seeding. Furthermore, the co-immunofluorescent analysis revealed that αSyn pathology preferentially affects specific neuronal subtypes, including mitral cells, inhibitory neurons (ChAT-positive), and calcium-binding protein-expressing interneurons (Calreticulin- and Parvalbumin-positive) (**Supplementary Figure 5**). This differential susceptibility is consistent with postmortem human studies (Beach et al., 2009a, 2009c; Ubeda-Bañon et al., 2010) and further suggests that the earliest consequences of αSyn accumulation may be a functional disruption within vulnerable neuronal circuits, rather than widespread cytotoxicity.

Most importantly, our data consistently point to neuroinflammation as the most robust and early pathological correlate of the αSyn pSer129 spreading. Gene expression analysis revealed a significant increase in pro-inflammatory markers, notably *Il-1β* and *Il-6* primarily in the OB and AON, that may be suggestive of NLRP3 inflammasome activation. Activation of the NLRP3 inflammasome pathway plays a critical role in neuroinflammation underlying PD pathogenesis (Jewell et al., 2022). NLRP3 inflammasome directly processes IL-1β into its active form (de Rivero Vaccari et al., 2014), whereas in some contexts IL-6 can induce NLRP3 inflammasome activation (Wang et al., 2020; Xiao et al., 2022). This inflammatory response was paralleled by quantitative histopathology showing a rapid and widespread glial activation across multiple olfactory structures. Specifically, we observed two distinct, yet overlapping, glial responses from microglia and astrocytes. In the case of microglial reactivity, αSyn PFF injection induced morphological changes (decrease in Area:Perimeter ratio) and microgliosis (increased cell number) in the OB, AON, APCX, and PCX. This microglial activation occurred as early as 1 month and was more pronounced at the earlier 1- and 2-month time points. The bilateral presence of reactive microglia in regions like the PCX further highlights a rapid, widespread response to the pathological αSyn spread, potentially contributing to the early, bilateral nature of olfactory dysfunction. In the case of astrocytes, astrogliosis (measured by increased GFAP-positive area and cell counts) also began early (1 month) in the aPCX and pPCX and was notably persistent at 3 months post-injection in the AON and aPCX. The persistence of astrogliosis suggests a sustained, rather than transient, inflammatory environment. The co-staining experiments demonstrated that αSyn pSer129 is not present within the GFAP-positive astrocytes, indicating that the astrocytes are likely reacting to paracrine signals released by αSyn-affected neurons or reactive microglia, possibly through pro-inflammatory pathways as suggested by the gene expression results. Other alternatives for the absence of αSyn pSer129 in astrocytes, despite prominent astrogliosis, may be inferred from recent clinical data (Altay et al., 2022) suggesting that astrocytic αSyn may undergo distinct post-translational modifications (such as nitration) not captured by standard pSer129 antibodies.

Given that our results suggest that inflammation may be a key driver for the previously reported functional impairment during this prodromal window, this model is uniquely positioned for testing anti-inflammatory interventions, including NLRP3 inhibitors, some of them already in early clinical trials for PD (Clarke et al., 2025; Luque et al., 2026), and other approaches such as neutralizing IL-1β (Dinarello, 2011) or anti-IL-6 (Jiao and Guo, 2022; Narazaki and Kishimoto, 2022), to determine if early modulation can improve functional impairment and arrest the progression toward neurodegeneration.

Taken together, these results support a model for prodromal PD in which the functional deterioration of the olfactory system is an initial consequence of αSyn spreading, which rapidly triggers a cascading neuroinflammatory response mediated by reactive microglia and persistent astrogliosis. Neuronal loss appears to be a delayed consequence, following a chronic period of unresolved inflammation. Defining the precise sequence of events, starting with αSyn seeding, then glial reactivity initiated by microglial activation and astrogliosis, leading to neuronal dysfunction with the final neuronal loss, is critical for identifying therapeutic windows that precede irreversible neurodegeneration. Future studies should focus on selectively modulating these early inflammatory responses to determine their causal link to the olfactory functional deficits.

## Conclusions

This study demonstrates that the injection of αSyn PFF into the olfactory bulb of wild-type mice results in the progressive, transneuronal spread of αSyn pSer129 pathology, which is accompanied by a robust and widespread neuroinflammatory response. Crucially, this neuroinflammation, characterized by increased microglial reactivity and persistent astrogliosis, is an early pathological process that precedes the onset of detectable neuronal loss in the AON and PCX.

The identification of a ‘neuroinflammation-only’ window, where pathology and functional deficits exist without neuronal loss, presents a therapeutic opportunity. This model provides a robust platform for testing repurposed anti-inflammatory agents to determine if early intervention can stabilize olfactory function and, crucially, halt the progression toward irreversible neurodegeneration.

## CRediT authorship contribution statement

**Gabriela Mercado:** Conceptualization, Methodology, Investigation, Formal analysis, Writing – original draft. **Lucas Stetzik:** Methodology, Investigation, Formal analysis. **Emily Kuhn:** Investigation. **Lindsay Meyerdirk:** Investigation. **Allison Lindquist:** Investigation. **Lorna Cohen:** Investigation, Resources (Van Andel Institute Optical Imaging Core, RRID:SCR_021968). **Jennifer A. Steiner:** Project administration, Supervision. **Martha L. Escobar Galvis:** Funding acquisition. **Darren Moore:** Project administration, Supervision, Resources. **Daniel Wesson:** Conceptualization, Funding acquisition, Supervision, Writing – review & editing. **Patrik Brundin:** Conceptualization, Funding acquisition, Supervision, Writing – review & editing.

## Declaration of Generative AI and AI-assisted technologies in the writing process

During the preparation of this work, the authors used Gemini 3 Flash (Google) to assist with grammar, phrasing and refining clarity. While using this AI-based tool, the authors carefully reviewed and edited the content as needed and take full responsibility for the content of the publication.

## Data Availability

The raw data supporting the conclusions of this article, including the high-resolution digitized whole-slide images and the AI model training parameters, will be made available by the authors, without undue reservation. Quantitative datasets are available from the corresponding author upon reasonable request.

## Acknowledgements

We thank the Van Andel Institute Vivarium (RRID:SCR_023211) for caring for the animals used in this study. The mouse PFFs used in the study were kindly provided by Dr. Kelvin Luk, University of Pennsylvania Perelman School of Medicine, USA. We thank Corinne Esquibel for her assistance with images acquisition (Van Andel Institute Optical Imaging Core, RRID:SCR_021968). We thank Lindsey Smith for her assistance with the development of AI models on the Aiforia® platform.

## List of Abbreviations

PD: Parkinson’s disease
αSyn: α-synuclein
pSer129: αSyn phosphorylated at Serine 129
PFF: recombinant αSyn preformed fibrils
AON: anterior olfactory nucleus
OB: olfactory bulb
PCX: piriform cortex
aPCX: anterior piriform cortex
pPCX: posterior piriform cortex
NeuN: Neuronal nuclear antigen
GFAP: Glial fibrillary acidic protein
Iba1: Ionized calcium-binding adaptor molecule 1

