## Supplementary Figures for "Neuroinflammation is an early pathological process in a mouse model of prodromal Parkinson’s disease"

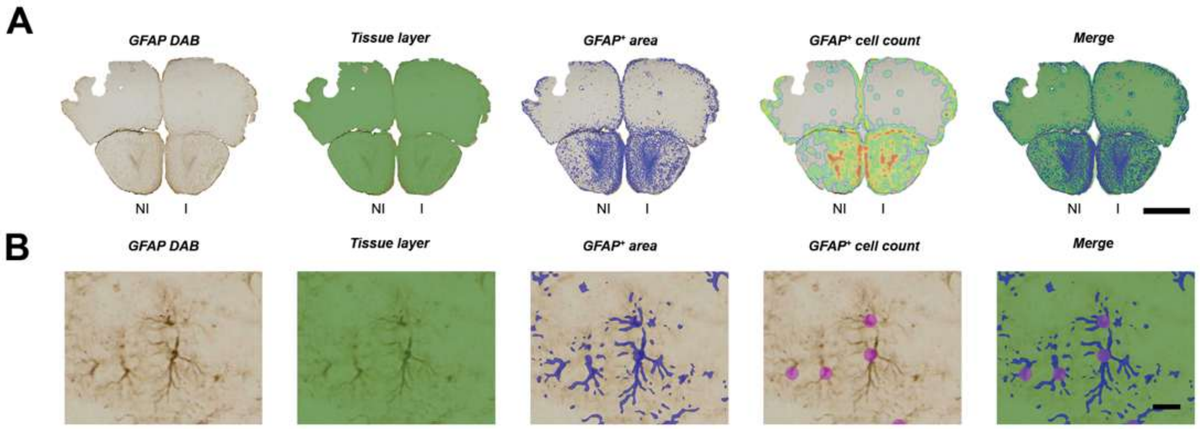


**Supplementary Figure 1. Validation of the deep learning model for automated astroglia quantification.** Representative visualization of the convolutional neural network (CNN) performance on an anti-GFAP DAB-stained coronal section of the AON (3 months post PFF-injection). **(A)** Sequential illustration of the multi-layered analysis: raw IHC image, automated tissue detection, semantic segmentation of GFAP-positive area, and individual soma detection (scale bar 1 mm). **(B)** High magnification inset demonstrating the model's ability to resolve individual astrocytic profiles from background noise (scale bar 20 μm). Non-injected (NI) and injected hemispheres (I), respectively.

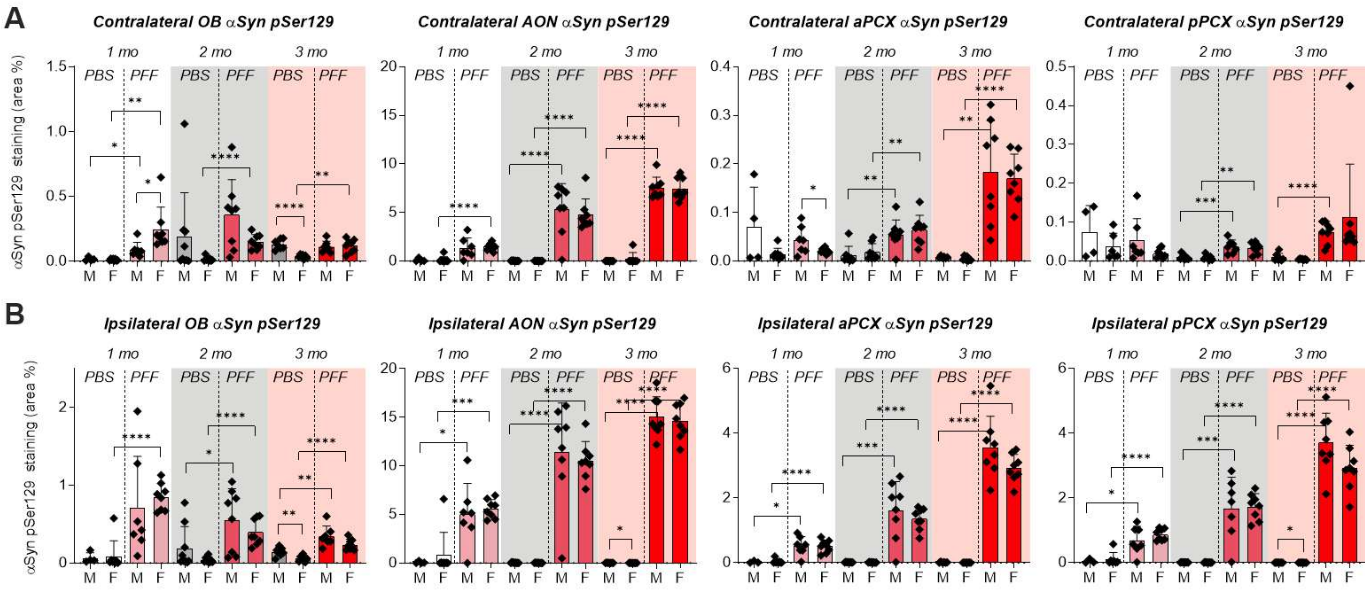


**Supplementary Figure 2. Dimorphic patterns in the contralateral propagation of αSyn pSer129 pathology.** Quantification of αSyn pSer129 burden in **(A)** contralateral and **(B)** ipsilateral olfactory structures of male (M) and female (F) mice. AI-assisted analysis reveals distinct sex-driven tendencies in the rate of inter-hemispheric spreading, particularly in the OB and aPCX at 1-month post-injection. Each data point represents the average per animal (n= 8 per group). Data were analyzed using three-way ANOVA (Factors: Time, Treatment, and Sex). Post-hoc comparisons between M and F, and treatment groups were performed using t-tests with a Bonferroni correction for multiple comparisons across time points.

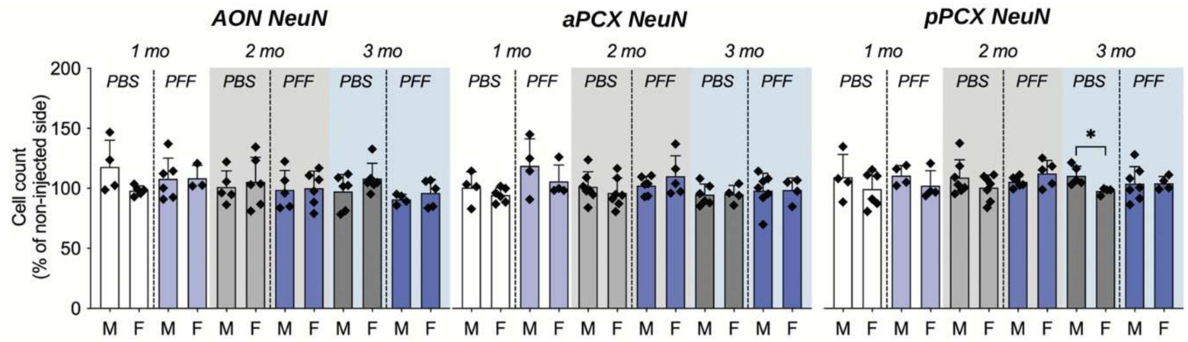


**Supplementary Figure 3. Stability of neuronal counts across sex cohorts.** Stereological quantification of NeuN-positive cells in male and female mice during the progressive spreading of αSyn pathology. Cell counts are expressed as a percentage of the non-injected (NI) hemisphere. Each data point represents the average per animal (n= 8 per group). Data were analyzed using three-way ANOVA (Factors: Time, Treatment, and Sex). Post-hoc comparisons between M and F, and treatment groups were performed using t-tests with a Bonferroni correction for multiple comparisons across time points.

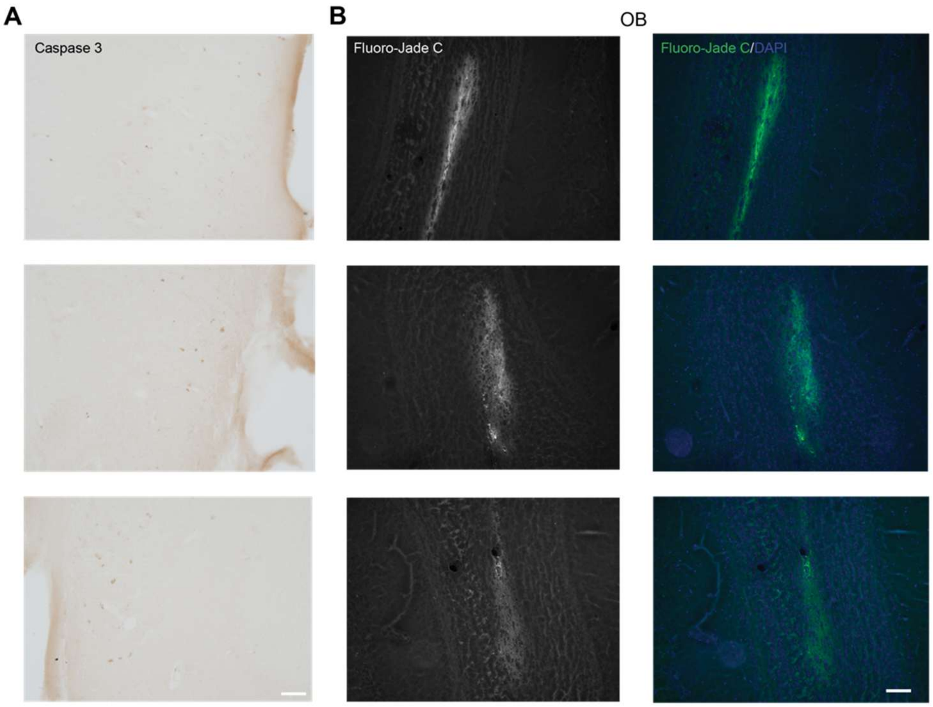


**Supplementary Figure 4. Assessment of acute cellular distress at injection-site 1 month post injections. (A)** Localized Caspase-3 immunoreactivity (apoptotic marker) within the dorsal olfactory nerve layer. **(B)** Fluoro-Jade C staining (green) restricted to the internal plexiform layer, corresponding to the mechanical needle track from stereotaxic delivery. These discrete signals were observed independently of αSyn PFF treatment, indicating they are surgical artifacts rather than pathology-driven neurodegeneration (scale bar: 50 μm).

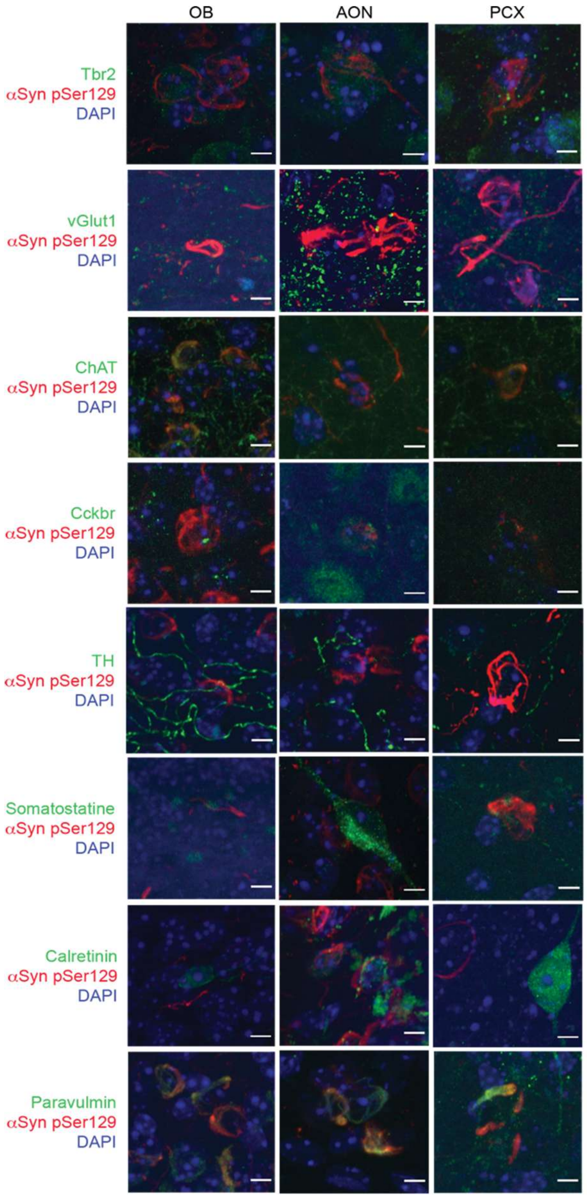


**Supplementary Figure 5. Phenotypic characterization of αSyn pSer129-positive neuronal subpopulations.** Representative confocal micrographs displaying the co-localization of αSyn pSer129 (red) with specific neurochemical markers (green) in the OB, AON, and PCX. Target populations include: Tbr2+ (mitral cells), vGlut1+ (glutamatergic), ChAT+ (cholinergic), Cckbr+ (interneurons), Somatostatin+, and the calcium-binding proteins Calretinin and Paravalbumin. Notably, TH+ (dopaminergic) interneurons were largely devoid of pathological inclusions. (DAPI counterstain in blue; scale bar: 5 μm).

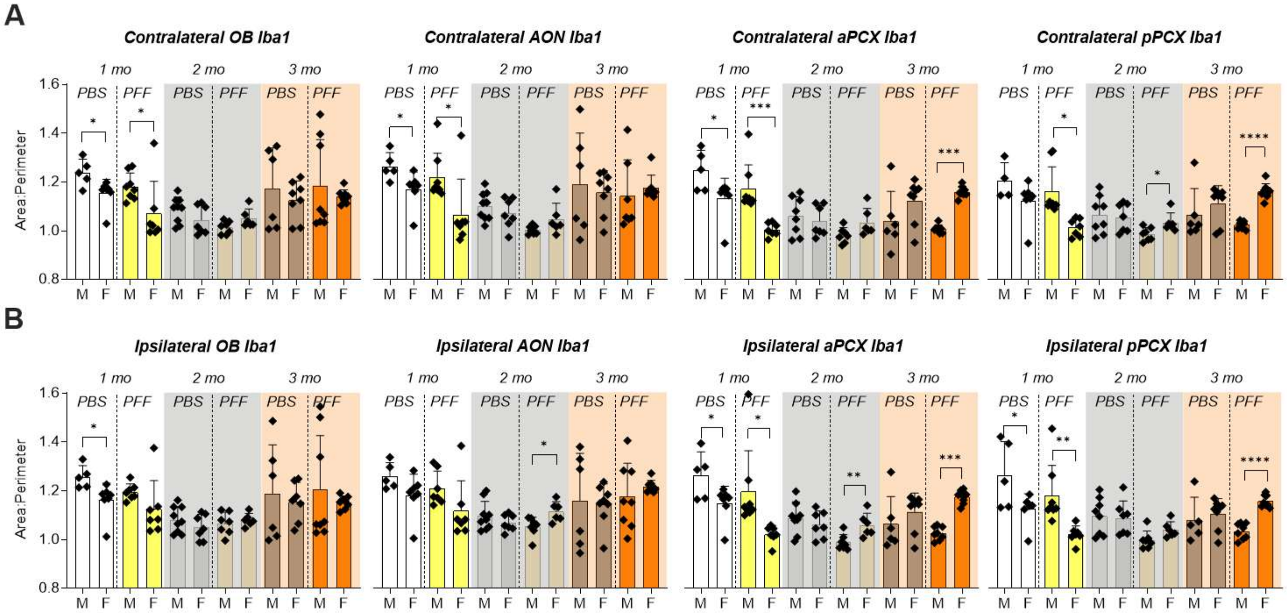


**Supplementary Figure 6. Sex-dependent variance in microglial reactivity.** AI-quantification of the Iba1+ Area:Perimeter ratio (microglial reactivity) and microgliosis (cell density) in **(A)** contralateral and **(B)** ipsilateral structures. Females displayed heightened microglial reactivity at 1 month, whereas males exhibited a delayed but significant response at 2 and 3 months exclusively in PFF-treated groups. Each data point represents the average per animal (n= 8 per group). Data were analyzed using three-way ANOVA (Factors: Time, Treatment, and Sex). Post-hoc comparisons between M and F were performed using t-tests with a Bonferroni correction for multiple comparisons across time points.

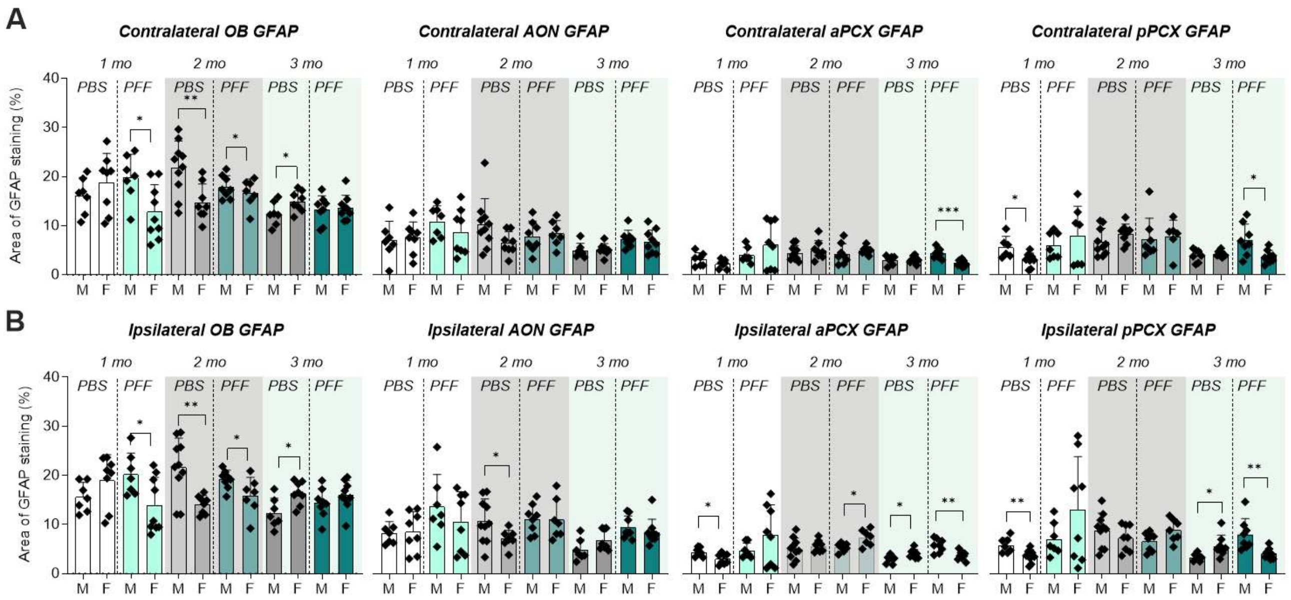


**Supplementary Figure 7. Sex-dependent variance in astrogliosis.** Sex-specific quantification of GFAP+ area percentage in the contralateral and ipsilateral hemispheres. Males exhibited increased baseline astrogliosis in specific regions. **(A)** contralateral (non-injected) and **(B)** ipsilateral (injected) hemisphere of the OB, AON, aPCX and pPCX of male (M) and female (F) mice. Each data point represents the average per animal (n= 8 per group). Data were analyzed using three-way ANOVA (Factors: Time, Treatment, and Sex). Post-hoc comparisons between M and F were performed using t-tests with a Bonferroni correction for multiple comparisons across time points.

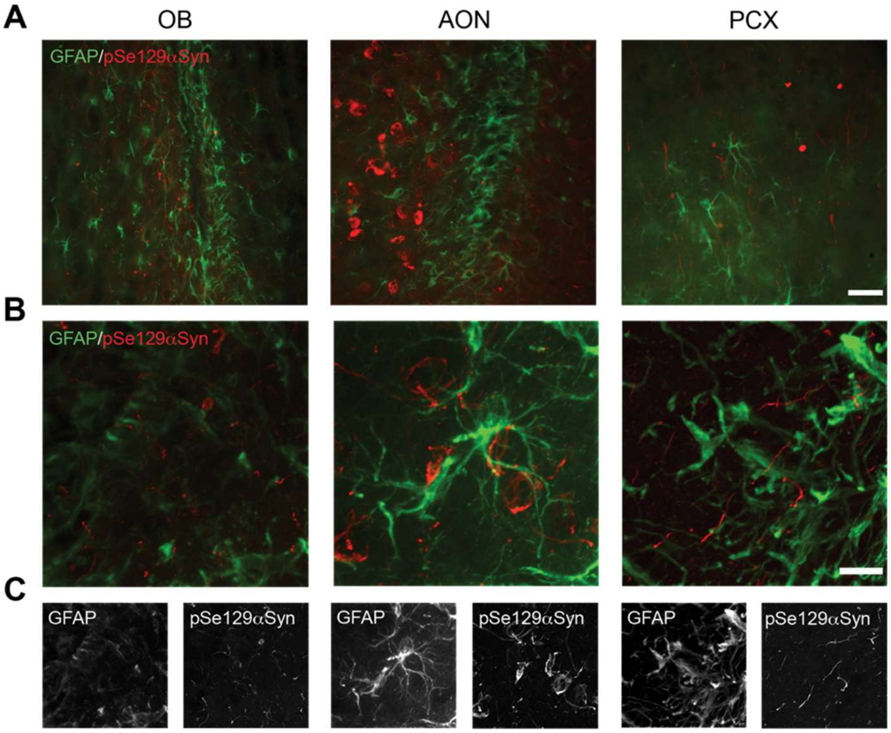


**Supplementary Figure 8. Astroglia αSyn pSer129 pathology interaction. (A)** Representative immunofluorescent images showing the spatial proximity of GFAP+ astrocytes (green) to αSyn pSer129+ neuronal inclusions (red). **(B)** High-magnification and **(C)** split-channel images demonstrate the absence of αSyn pSer129 within the astrocytic cytoplasm (scale bars: 250 μm and 10 μm).

**Supplementary Tables**

**Supplementary Table 1. TaqMan® probes used for real time PCR in the study.**

| Protein | Gene | Probe ID |
|---|---|---|
| α-synuclein | *Snac* | Mm01188700_m1 |
| Psd95 | *Dlg4* | Mm00492193_m1 |
| Chop | *Ddit3* | Mm01135937_g1 |
| Nlrp3 | *Nlrp3* | Mm00840904_m1 |
| Il-1β | *Il-1β* | Mm00434228_m1 |
| Gfap | *Gfap* | Mm01253033_m1 |
| Il-6 | *Il-6* | Mm00446190_m1 |
| Bip | *Hspa5* | Mm00517691_m1 |
| Nox2 | *Cybb* | Mm01287743_m1 |
| Gapdh | *Gapdh* | Mm99999915_g1 |

**Supplementary Table 2. Antibodies used for IHC staining in the study.**

| Antibody | Concentration | Supplier | Catalog number |
|---|---|---|---|
| *Primary antibodies IHC* | | | |
| Mouse anti-NeuN | 1:1000 | Millipore | Ab377 |
| Rabbit anti-pSer129 αSyn | 1:1000 | Abcam | Ab51253 (EP1536Y) |
| Rabbit anti-GFAP | 1:1000 | Abcam | Ab7260 |
| Rabbit anti-Iba1 | 1:1000 | Wako | 019-19741 |
| Rabbit anti-Active-Caspase 3 | 1:1000 | BD Pharmagen | 559565 |
| *Secondary antibodies IHC* | | | |
| Goat anti-mouse Biotinylated | 1:500 | Vector Laboratories | BA-9200-1.5 |
| Goat anti-rabbit Biotinylated | 1:500 | Vector Laboratories | BA-1000-1.5 |

**Supplementary Table 3. Antibodies used for IF staining in the study.**

| Antibody | Concentration | Conditions | Supplier | Catalog/Clone number |
|---|---|---|---|---|
| *Primary antibodies IF* | | | | |
| Rabbit anti-pSer129 αSyn | 1:1000 | 4˚C o/n | Abcam | Ab51253 (EP1536Y) |
| Mouse anti-pSer129 αSyn | 1:1000 | 4˚C o/n | Abcam | Ab184674 |
| Mouse anti-Calretinin | 1:100 | HIER RT o/n | Swant | 6B3 |
| Rat anti-Somatostatin | 1:50 | HIER RT o/n | Abcam | ab30788 |
| Rabbit anti-Parvalbumin | 1:500 | 4˚C o/n | Abcam | ab181086 |
| Chicken anti-Tbr2 | 1:50 | HIER RT o/n | Millipore | AB15894 |
| Goat anti-Cckbr | 1:500 | 4˚C o/n | Thermo Scientific | PA5-18384 |
| Rabbit anti-TH | 1:1000 | RT o/n | Millipore | 657012 |
| Rabbit anti-vGlut1 | 1:2000 | 4˚C o/n | Synaptic Systems | 135 303 |
| Rabbit anti-ChAT | 1:500 | 4˚C o/n | Abcam | ab178850 |
| Rabbit anti-GFAP | 1:2000 | 4˚C o/n | Abcam | ab7260 |
| *Secondary antibodies IF* | | | | |
| Donkey anti-chicken FITC | 1:1000 | 2 h RT | ThermoFisher | SA1-72000 |
| Donkey anti-goat Alexa-488 | 1:1000 | 2 h RT | ThermoFisher | A-11055 |
| Donkey anti-rabbit Alexa-488 | 1:1000 | 2 h RT | ThermoFisher | A-21206 |
| Donkey anti-mouse Alexa-488 | 1:1000 | 2 h RT | ThermoFisher | A-21202 |
| Donkey anti-rat Alexa-488 | 1:1000 | 2 h RT | ThermoFisher | A-21208 |
| Donkey anti-rabbit Alexa-555 | 1:1000 | 2 h RT | ThermoFisher | A-31572 |
| Donkey anti-mouse Alexa-555 | 1:1000 | 2 h RT | ThermoFisher | A-31570 |
